# Darwin: A Hardware-acceleration Framework for Genomic Sequence Alignment

**DOI:** 10.1101/092171

**Authors:** Yatish Turakhia, Kevin Jie Zheng, Gill Bejerano, William J. Dally

## Abstract

Genomics is set to transform medicine and our understanding of life in fundamental ways. But the growth in genomics data has been overwhelming - far outpacing Moore’s Law. The advent of third generation sequencing technologies is providing new insights into genomic contribution to diseases with complex mutation events, but have prohibitively high computational costs. Over 1,300 CPU hours are required to align reads from a 54× coverage of the human genome to a reference (estimated using [1]), and over 15,600 CPU hours to assemble the reads *de novo* [2]. This paper proposes “Darwin” - a hardware-accelerated framework for genomic sequence alignment that, without sacrificing sensitivity, provides 125× and 15.6× speedup over the state-of-the-art software counterparts for reference-guided and *de novo* assembly of third generation sequencing reads, respectively. For pairwise alignment of sequences, Darwin is over 39,000× more energy-efficient than software. Darwin uses (i) a novel filtration strategy, called D-SOFT, to reduce the search space for sequence alignment at high speed, and (ii) a hardware-accelerated version of GACT, a novel algorithm to generate near-optimal alignments of arbitrarily long genomic sequences using constant memory for trace-back. Darwin is adaptable, with tunable speed and sensitivity to match emerging sequencing technologies and to meet the requirements of genomic applications beyond read assembly.

## 1. INTRODUCTION

Since the completion of the first draft of the human genome in 2001 [3, 4], genomic data has been doubling every 7 months - far outpacing Moore’s Law. By 2020, genomic data is expected to reach exabyte scale and surpass Youtube and Twitter by 2025 [5]. This growth has been primarily led by the massive improvements in “genome sequencing” - a technology that enables reading sequence of nucleotide bases from DNA molecules. Today, it is possible to sequence genomes on rack-size, high-throughput machines at nearly 50 human genomes per day [6], or on portable, USB-stick size sequencers that require several days per human genome [7]. This data explosion has been immensely valuable to the emerging field of personalized medicine [8] and in detecting when some genomic mutations predispose humans to certain diseases, including cancer [9], autism [10] and aging [11]. It has also contributed to the understanding of the molecular factors leading to phenotypic diversity (such as eye and skin color, etc.) in humans [12], their ancestry [13] and the genomic changes that made humans different from related species [14].

Third generation technologies produce much longer *reads* of contiguous DNA — tens of kilobases compared to only a few hundred bases in both first and second generation [15]. This drastically improves the quality of the *genome assembly* — where the reads are assembled into a complete sequence. For example, contiguity in the Gorilla genome assembly was recently found to improve by over 800× using long reads [16]. For personalized medicine, long reads are superior in (i) identifying structural variants i.e. large insertions, deletions and re-arrangements in the genome spanning kilobases of more which are sometimes associated with diseases, (ii) haplotype phasing, to distinguish mutations on maternal versus paternal chromosomes and (iii) resolving highly repetitive regions in the genome [15].

Despite several advantages, the long read technology suffers from one major drawback - high error rate in sequencing. Mean error rates of 15% on Pacific Biosciences (PacBio) technology and up to 40% on Oxford Nanopore Technology (ONT) have been reported [17, 18]. These errors are corrected using computational methods that can be orders of magnitude slower than those used for the second generation technologies. For instance, using BWA [1], aligning 54× coverage reads of the human genome to reference requires over 1,300 CPU hours on PacBio reads (15% error rate), compared to only 270 CPU hours for short Illumina reads (*<*2% error rate).

In this paper, we describe a hardware-acceleration framework for genome sequence alignment, named *Darwin*, which accelerates the compute-intensive task of long-read assembly by orders of magnitude. We note that Darwin is a general framework for acceleration of sequence alignment, and can be used for applications beyond read assembly, such as homology search [19] or whole genome alignments [20], but we specifically focus on Darwin’s application to long read sequencing due to it’s far-reaching potential in near-term.

This paper makes the following contributions:

1. We propose <u>G</u>enome <u>A</u>lignment using <u>C</u>onstant-memory <u>T</u>race-back (GACT) - a novel algorithm using dynamic programming for aligning arbitrarily long sequences using constant memory for the compute-intensive step. This has a profound hardware design implication, as all previous hardware accelerators for genomic sequence alignment have assumed an upper-bound on the length of sequences they align (typically <10Kbp) [21, 22] or have left the trace-back step in alignment to software [23, 24], thus undermining the benefits of hardware-acceleration. We show that GACT can produce alignments which are within 0.95% of the optimal and have no effect on the consensus accuracy for coverage above 13×.
2. We propose <u>D</u>iagonal-band <u>S</u>eed <u>O</u>verlapping based Filtration Technique (D-SOFT) - a novel filtering algorithm to reduce the search-space for dynamic programming. D-SOFT can be used for finding read-to-reference mapping in reference-guided assembly or for finding overlaps between reads in *de novo* assembly at high speed and sensitivity. D-SOFT filters by counting the number of non-overlapping bases in a band of diagonals. This allows D-SOFT to be tuned flexibly — at only 0.16% sensitivity loss, D-SOFT can be made 121× more specific, thus alleviating the need for additional filtering heuristics.
3. We propose a hardware design for acceleration of GACT. The design is fully prototyped on FPGA. We perform ASIC synthesis and layout for GACT on 45nm TSMC technology node. With our ASIC implementation, pairwise alignment of sequences has a speedup of 762× over software and is over 39,000× more energy-efficient.
4. We propose Darwin, a framework which combines hardware-accelerated GACT with D-SOFT. We evaluate Darwin’s speedup in comparison to software counterparts and show that Darwin can be tuned to a wide-range of speed and sensitivity. Darwin provides a speedup of 125× and 107× for reference-guided assembly of the human genome using PacBio and ONT 2D reads, respectively, without losing sensitivity. For finding read overlaps for *de novo* assembly of PacBio reads, Darwin provides a 15.6× speedup over DALIGNER [2].

The rest of the paper is organized as follows. Section 2 provides relevant background in genome sequence alignment, sequencing and read assembly. Section 3 and 4 present the GACT and the D-SOFT algorithms, respectively. Section 5 explains how Darwin’s framework combines D-SOFT and hardware-accelerated GACT to perform reference-guided and *de novo* assembly of long reads. Section 6 provides the hardware design and implementation details to accelerate the GACT algorithm. We provide experimental methodology in Section 7, and discuss the results in Section 8. Section 9 discusses some relevant related work, which is followed by conclusion and future work discussion in Section 10.

## 2. BACKGROUND

In this section, we provide some necessary background on genome sequence alignment, genome sequencing and genome assembly needed for the rest of this paper.

### 2.1 Genome sequence alignment

The sequence alignment problem can be formulated as follows: Given a query sequence *Q* = *q*_1_, *q*_2_, .., *q*_*n*_ and a reference sequence *R* = *r*_1_, *r*_2_, .., *r*_*m*_ (*m ≥ n*), assign gaps (denoted by ‘-’) to *R* and *Q* to maximize an alignment score. The alignment assigns letters in *R* and *Q* to a single letter or a gap in the opposite sequence. For genomic sequences, the letters are in the alphabet Σ = {*A,C, G, T*}, corresponding to the four nucleotide bases.

Sequence alignment often uses the Smith-Waterman algorithm [25], since it provides optimal *local alignments* for biologically-meaningful scoring schemes. A typical scoring scheme rewards matching bases and penalizes mismatching bases, with these rewards and penalties specified by a substitution matrix *W* of size Σ × Σ. The gaps in *R* and *Q*, known as *insertions* and *deletions*, are penalized by another parameter, *gap*. Smith-Waterman operates in two phases: (i) matrix-fill, in which a score *H*(*i, j*) and a trace-back pointer is computed for every cell (*i, j*) of the DP-matrix having dimensions (*m* + 1) × (*n* + 1) using the dynamic programming equations of Figure 1a, and (ii) trace-back, which uses trace-back pointers, starting from the highest-scoring cell, to reconstruct the optimal *local* alignment. Figure 1c shows an example DP-matrix computed using the parameters in Figure 1b for sequences *R* =‘GGCGACTTT’ and *Q* =‘GGTCGTTT’. An arrow in each cell represents the trace-back pointer, which points to the cell that led to its score, with no arrow for cells with score 0, where the trace-back phase must terminate. Red arrows indicate that trace-back path from the highest-scoring cell (highlighted using red) to reconstruct the optimal alignment shown in Figure 1d.

**Figure 1:**
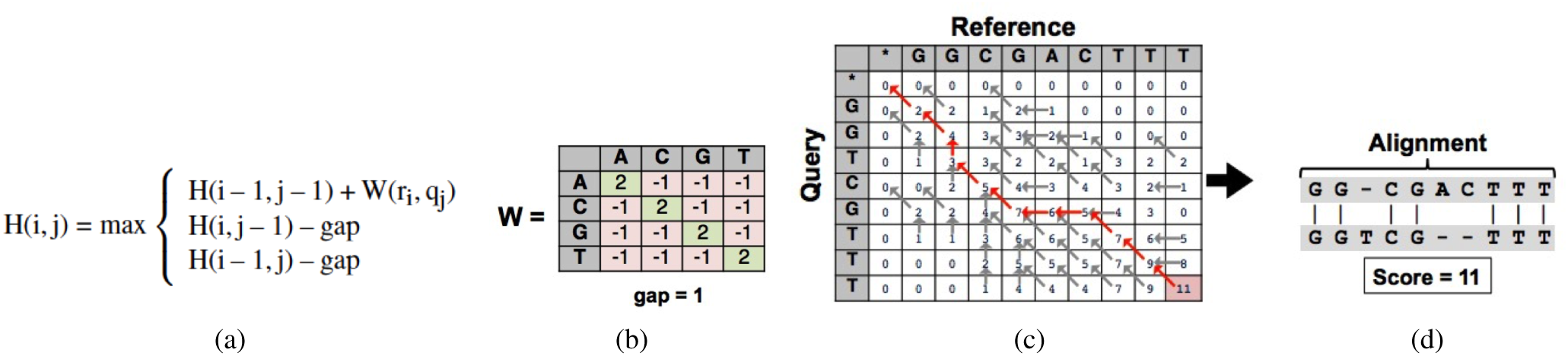
(a) Smith-Waterman equations. (b) Smith-Waterman scoring parameters. (c) DP-matrix with trace-back pointers. (d) Optimal alignment.

Smith-Waterman is a highly compute intensive algorithm with time complexity of 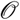(*mn*). To align even a 1000 bp read with the 3 × 10^9^ bp human genome has a complexity of 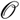(10^12^) and takes several minutes on highly-optimized software. As a result, most alignment heuristics use an additional filtration step based on the *seed-and-extend* paradigm, made popular by BLAST [19]. This approach uses substrings of fixed size *k* from *Q*, known as *seeds*, and finds their exact matches in *R*, known as *seed hits*. Finding seed hits is 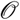(*m*), but can be achieved in 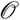(1) with some pre-processing on *R* [19]. Once the seed hits are found, the extend step uses dynamic programming equations to compute DP-matrix scores for only the cells surrounding the seed hit that could result in a good local alignment, thereby avoiding the high cost of computing every cell of the DP-matrix. Alignments between *R* and *Q* having no exactly matching substrings of size *k* or larger will not be discovered by this approach, resulting in lower *sensitivity*. The *seed-and-extend* paradigm trades sensitivity for speed but works well in practice and has been incorporated in a large number of alignment heuristics since BLAST [26, 20, 1, 27, 28, 29, 2], each heuristic optimized for a speed/sensitivity trade-off point depending on the application context and the properties of sequences in that context.

### 2.2 Genome sequencing and assembly

Genome/DNA sequencing is the process of shearing DNA molecules of the genome into a large number *N* of random fragments and reading the sequence of bases in each fragment. The size of the fragments depends on sequencing technology. Long read sequencing technologies, such as Pacific Biosciences and Oxford Nanopore, have a mean read length *L* of about 10Kbp. Typically, *N* is chosen such that each base-pair of the genome *G* is expected to be covered more than once, i.e. *C* = (*NL/G*) *>* 1, where *C* is known as the *coverage* of sequencing. Higher sequencing coverage has the advantage that each base-pair is likely covered by several reads and small errors in each read can be corrected using a *consensus* of multiple reads. Despite high error-rate of 15% at base-pair level, Pacific Biosciences reads have consensus accuracy of *>* 99.99% at a coverage of 50× [30].

Genome assembly is the computational process of reconstructing a genome *G* from a set of reads *S*. In *reference-guided* assembly a reference sequence *R* is used to assist the assembly. In *de novo* assembly, *G* is constructed only from *S*, without the benefit (or bias) of *R*. The preferred approach to *de novo* assembly of long reads is using *overlap-layout-consensus* (OLC) [31]. In the *overlap* phase, all read pairs having a significant overlap are found. During the *layout* phase, overlapping reads are merged to form larger contigs. Finally, in the *consensus* phase, the final DNA sequence is derived by taking a consensus of reads, which corrects the vast majority of read errors. By far, the most time-consuming step of OLC is the *overlap* step, which requires 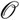(*N*^2^) time for *N* reads.

In reference-guided assembly, sometimes called *resequencing*, the reads are directly aligned to reference sequence *R*, which closely resembles the sequenced genome. Reference-guided assembly is 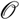(*N*) for *N* reads, produces fewer contigs compared to *de novo* assembly, and is good at finding small changes, called *variants*, of the sequenced genome with respect to the reference genome. However, it can also introduce reference-bias, and unlike *de novo* assembly, reference-guided assembly does not capture large *structural variants* between reference and sequenced genome. The debate on the relative merits of *de novo* and reference-guided assembly for clinical practice is far from settled, but the current standard of BWA-GATK pipeline [32], incorporates best of both worlds — BWA performs fast reference mapping, and GATK implements techniques to reduce reference-bias and identify *de novo* variants in the sequenced genome. Additionally, graph-based algorithms, such as Manta [33], are now being used for detecting structural variants for certain clinical applications.

## 3. GACT ALGORITHM

### Motivation

The advent of long-read technology has made the problem of efficiently aligning two long genomic sequences critical. Smith-Waterman requires 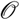(*mn*) space to optimally align two sequences of lengths *m* and *n*. Hirschberg’s algorithm [34] can improve the space complexity to linear 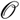(*m* + *n*), but is rarely used in practice because of its performance. As a result, linear-space heuristics, such as Banded Smith-Waterman [35] and X-drop [36], that do not guarantee optimal alignments but have linear space and time complexity, have become popular. For long reads, both algorithms require prohibitive trace-back pointer storage for hardware implementation. The longest recorded read is over 300Kbp [37], storing a band of 2-bit trace-back pointers of width *B* on both sides of the main diagonal requires 2*B* × 300 × 10^3^ × 2 bits, or roughly 15MB for *B* = 100. Not only is provisioning such a large memory using on-chip SRAM difficult, it is also performance- and energy-inefficient, and restricts the amount of on-chip parallelism. We later show in Section 8, that even a band *B* = 100 is insufficient for long reads since they have very different insertion and deletion rates, resulting in the optimal alignment drifting far from the diagonal.

### Description

GACT for left extension, i.e. aligning sequences *R* and *Q* to the left of position *i*^*∗*^ in *R* and *j*^*∗*^ in *Q*, is shown in Algorithm 1. GACT works in the *seed-and-extend* paradigm, and positions *i*^*∗*^ and *j*^*∗*^, where *R* and *Q* are likely to have an alignment, are generated by the seeding stage. GACT finds alignment between long sequences by incrementally finding optimal alignments between a series of smaller subsequences of *R* and *Q*, starting at (*i*^*∗*^, *j*^*∗*^). The size of the subsequences have an upper-bound of tile size *T*, an input parameter to GACT. GACT maintains the start and end positions of *R* (*Q*) in the current tile in *i*_*start*_ ( *j*_*start*_) and *i*_*curr*_ ( *j*_*curr*_), respectively.

The function *Align* computes the optimal alignment for the current tile. The alignment begins from the highest-scoring cell of the DP-matrix if argument *t* is true. Otherwise the alignment starts from the bottom-right cell. Aligning from highest-scoring cell is used when the filtering stage generates approximate (*i*^*∗*^, *j*^*∗*^) around which alignment should begin, and is used only for the first tile in GACT. Aligning from bottom-right cell is used to “stitch” together the alignments from successive tiles during extension. *Align* returns (i) the alignment score of the tile *TS*, (ii) the offsets *i*_*off*_ and *j*_*off*_ in *R* and *Q* based on the number of bases of each sequence participating in the alignment, (iii) a set of traceback pointers *tb*, that can be used to used to reconstruct the final alignment, and (iv) the position of the highest-scoring cell (*i*_*max*_, *j*_*max*_) if *t* is set. GACT terminates when it reaches the start of *R* or *Q*, or when the trace-back results in less than half of the bases in *R* and *Q* being aligned, indicating that further extension is unlikely to result in a positive score. For right extension, GACT works similar to Algorithm 1, but using reverse of *R* and *Q*. Because the maximum length of sequences used as input for *Align* is *T*, a maximum of *T*^2^ trace-back pointers are stored in memory. The size of the traceback buffer, *tb*_*le f t*, which stores global trace-back for left extension, is still linear in sequence length (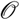(*m* + *n*)), but the compute-intensive step of *Align* is implemented in constant memory (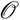(*T*^2^)).

#### Algorithm 1

GACT for Left Extension

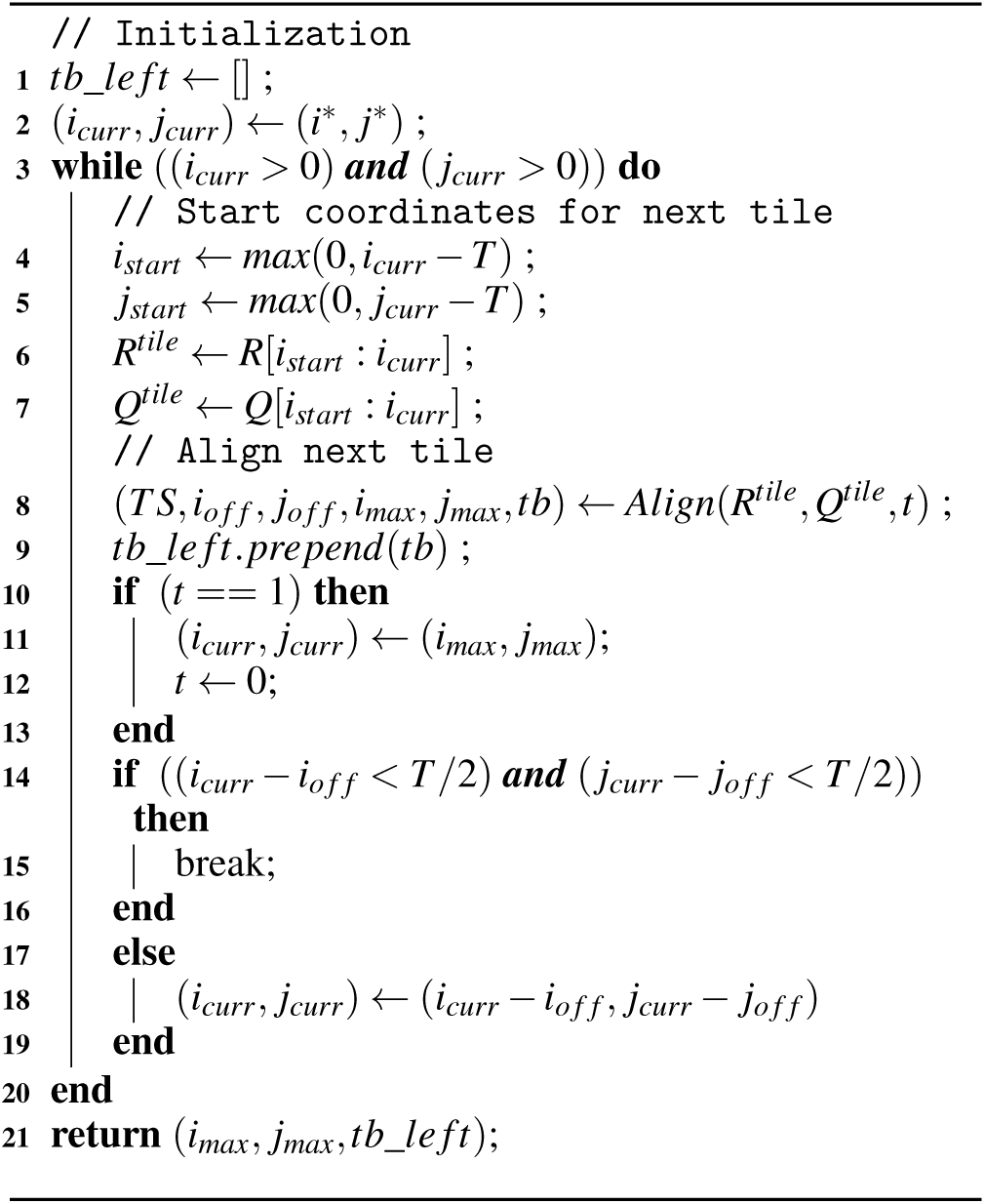

Figure 2 provides an example of GACT using *T* = 5, for the two sequences of Figure 1. The figure also illustrate why

**Figure 2:**
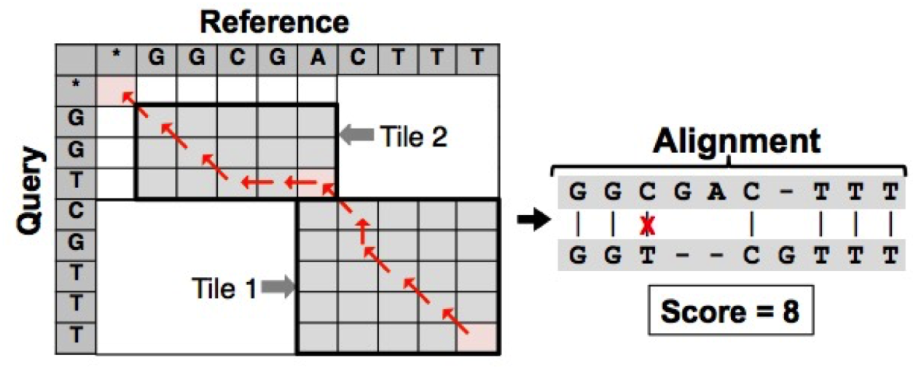
GACT tiling and alignment with sequences and scoring scheme of Figure 1 and *T*=5. Gray cells represent computed cells (scores not shown) and red path represents GACT trace-back.

GACT is not optimal - the resulting alignment is different from the optimal alignment of Figure 1d, and with a lower alignment score. This is because GACT only computes scores of the gray and red cells of a single tile of the DP-matrix of Figure 2 at any point of time, leaving out the white cells. So while GACT constructs *optimal* alignments from the starting cell (*i*_*curr*_, *j*_*curr*_) within a tile, it cannot guarantee the resulting alignment to be *globally* optimal once the alignments from individual tiles are “stitched” together. We empirically show in Section 8, that for *T* = 300, GACT produces alignments that are within 0.95% (worst-case) of the optimal score for sequences with 70% similarity.

GACT has a number of unique features: (i) its performance is linear (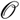(*max*(*m, n*) · *T*)) in the length of the sequences being aligned, as opposed to quadratic for optimal alignment, (ii) its compute-intensive step *Align* has a constant trace-back memory requirement, 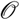(*T*^2^), that depends only on the tile size *T*, and not on the length of sequences being aligned, which makes it suitable for hardware implementation, (iii) unlike banded alignment, which imposes a strict band around the main diagonal, GACT’s band is adaptive, as it is adjusted depending on the trace-back path it has encountered so far. (iii) is an extremely useful feature for long reads, that have different insertion and deletion rates - resulting in the alignment drifting away from the main diagonal.

## 4. D-SOFT: A FILTERING ALGORITHM FOR SEQUENCE ALIGNMENT

### Motivation

Simple *seed-and-extend* algorithms, such as BLAST, can result in excessive computation due to a high rate of false positives [38]. For a seed of size *k* bp, an error rate of *e*, and uniformly distributed bases, *G/*4^*k*^ false positives are expected, while the probability of a true positive is (1 − *e*)^*k*^. For *G* = 3 × 10^9^, *e* = 0.3, and *k* = 12, 178 false positives are expected but only 1.38% of the actual matches will be detected. To address this issue, Long read assembly algorithms, such as [29, 39], use a generalization of the two-hit BLAST proposed in [38], and filter by using multiple seeds from the read, counting the number of matching seeds in a band of diagonals. A match is detected when the count exceeds a threshold, *d*. For high sensitivity, several closely-spaced seeds from the query sequence are used, as they have a higher probability of staying within a narrow diagonal band. This method has high sensitivity but poor specificity, as noted in [2]. A large number of false positives occur when overlapping seeds contribute to the count. For example, if every seed from the query is used (i.e. *stride* = 1), only *k* + *d* consecutive bases must match exactly for a region to be chosen. [2] improves on this technique by counting the actual number of non-overlapping bases (instead of the number of seeds) in a band of diagonals and then filtering on a fixed threshold. This results in better specificity and speedup, since fewer additional filter stages are required. Here, we propose D-SOFT, that like [2] counts the number of unique matching bases in a diagonal, but differs in its implementation strategy. We highlight the advantages of D-SOFT versus [2] later in Section 9.

### Description

Like [2], D-SOFT filters by counting the number of unique bases in *Q* covered by seed hits in diagonal bands. Bands in which the count exceeds *h* bases are selected for alignment. Figure 3 illustrates the D-SOFT technique. The X- and Y-axes in the figure correspond to positions in *R* and *Q*, respectively. Similar to [2], *R* is divided up into *N*_*B*_ *bins*, of size 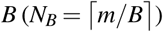. In Figure 3, *N*_*B*_ = 6. Each bin is associated with one uniquely colored diagonal band, with diagonals having a slope of 1. Ten seeds with *k*=4 from *Q* are used from consecutive positions, starting at position 1. The seed start positions are marked by dotted horizontal lines. For each seed, if a hit is found in *R*, a red dot is marked on its horizontal line, with its X-coordinate equal to the hit position in *R*. The tail of the red dot covers *k*=4 bases, showing the extent of the match. The count over each diagonal band is equal to the number of bases in *Q* covered by seed hits in that band. For instance, seeds at positions 4, 5 and 6 in *Q* find a hit in bin 1 of *R*, but its count is 6 since the seed hits cover only 6 bases (4-9) in *Q*. Bin 3 also has 3 seed hits, but the hits cover 9 bases (1-4 and 5-9). Similarly, the count of bin 4 is 4 with two seed hits, since both hits result from seed at query position 3 covering same bases (3-6). If *h* = 8 is used, only bin 4 would be chosen as a candidate. However, if a seed hit count strategy is used, both bin 1 and bin 3 would have the same count (=3), even though bin 3, having more matching bases, has a better probability of finding a high-scoring alignment. Counting unique bases in a diagonal band allows D-SOFT to be more specific than strategies that count seed hits, because overlapping bases are not counted multiple times.

**Figure 3:**
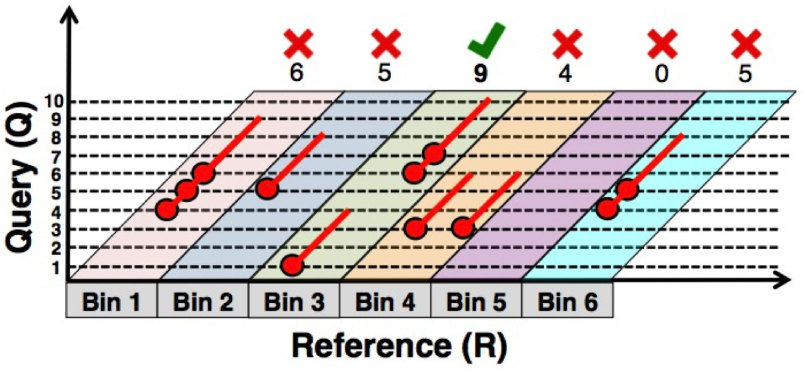
Illustration of D-SOFT algorithm for *k*=4, *N*=10, *h*=8, *N*_*B*_=6.

As shown in Algorithm 2 D-SOFT operates using two arrays, *last*_*hit*_*pos* and *bp*_*count*, each of size *N*_*B*_ to store the last seed hit position for every bin and the total bases covered by seed hits, respectively. Seeds of size *k* are drawn from *Q*, with a specified *stride* between *start* and *end* (lines 4-5). For each seed, function *SeedLookup* finds all hits in *R* (line 6). For each *hit*, its *bin* and its *overlap* with the last hit in the bin is computed (lines 7-9). Non-overlapping bases are added to *bp*_*count* (line 11). When the *bp*_*count* for a new bin exceeds the threshold *h*, the last hit in the bin is added to a filtered list of candidate positions, *candidate*_*pos* (lines 12-13). This list is returned at the end of the algorithm (line 17) and is used as input to the GACT algorithm. In algorithm 2, we use unspaced seeds separated by a fixed stride, but D-SOFT can be easily generalized to spaced seeds [40], or even non-strided seed selection using more involved techniques [41], which can help further improve the sensitivity. Moreover, D-SOFT can work with any implementation of *SeedLookup*, such as seed position table [19, 20], suffix arrays or trees [42], or compressed index tables based on Burrows Wheeler Transform [27], FM-index [43], etc.

#### Algorithm 2

D-SOFT

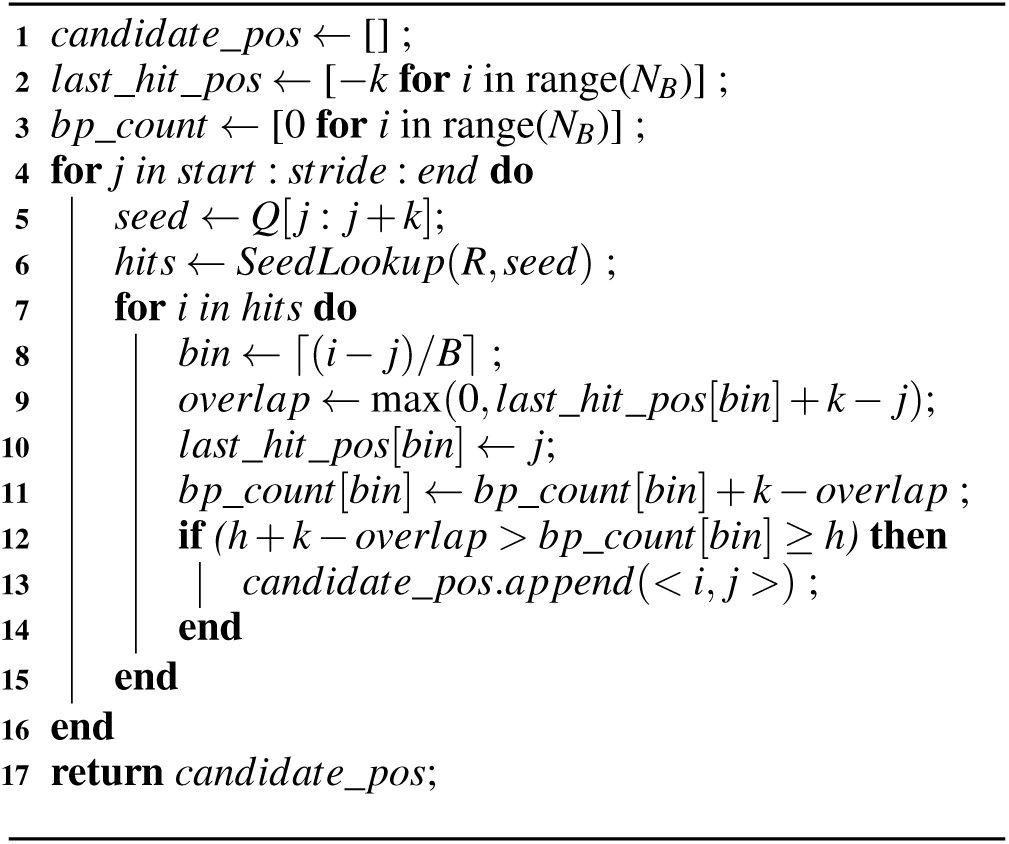

D-SOFT can be used for reference-guided as well as *de novo* assembly of long reads. In reference-guided assembly, the reference genome is used as *R* in Algorithm 2, and the reads are processed one at a time, using their forward as well as reverse-complement sequences, as *Q*.

For *de novo* assembly, Figure 4 shows how D-SOFT can be used to infer overlaps between reads. The *P* reads *r*_1_, *r*_2_, .., *r*_*P*_ are concatenated together to form *R*. The reads are made to align with the bin boundaries, filling the space between boundaries using unknown nucleotides, ‘N’. This way, each bin is associated with a unique read and this information is stored in a dictionary. Then, the forward and reverse-complement of each read *r*_*i*_ is used as *Q* to infer overlaps with other reads, as shown in Figure 4. When *P* is very large, exceeding the memory capacity, the reads are partitioned into blocks that fit into available memory. For each block, a different *R* is constructed.

**Figure 4:**
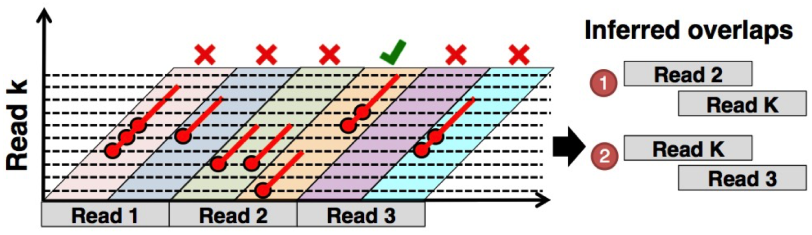
Inferring read overlaps using D-SOFT.

## 5. DARWIN: SYSTEM OVERVIEW

Darwin is a general-purpose framework for genomic sequence alignment that combines hardware-accelerated GACT with D-SOFT. Its design is based on two observations: (i) the parameters of D-SOFT can be set to make it very specific even for noisy sequences at high sensitivity, and (ii) the hardware-acceleration of GACT results in 762× speedup over software. This allows Darwin to directly use dynamic programming in GACT following filtering of D-SOFT and still provide high speedup. Using a minimum threshold, *h*_*tile*_, on the score of the first tile in GACT (line 8, Algorithm 1), avoids processing further tiles for most false positives. Further filtering of D-SOFT’s false positives can be handled by rejecting all D-SOFT candidates that result in shorter than expected extensions using GACT. This is explained in section 7. For simplicity, Darwin uses *B* = 100, *stride* = 1 and *start* = 0 in D-SOFT, reducing the number of D-SOFT parameters to (*k, N, h*), where *N* = *end* in Algorithm 2. GACT uses tile size, *T* and the minimum score threshold of the first tile, *h*_*tile*_, as the two parameters.

For high performance, Darwin uses an uncompressed seed position table similar to LASTZ [20] for *SeedLookup* in D-SOFT. To reduce excessive false positives from repetitive regions, we exclude high frequency seeds i.e. seeds occurring more than 32 times *G/*4^*k*^, the expected frequency of the seed hit for a uniform random genome sequence of size *G*. Darwin uses 16-bit integers for *last*_*hit*_*pos* and *bp*_*count*, requiring 120MB for human genome and *B*=100.

## 6. HARDWARE DESIGN AND IMPLEMENTATION OF GACT

In this section, we present the design of the *GACT array*, custom hardware that accelerates the compute-intensive *Align* routine in GACT, as described in Section 3. The rest of Algorithm 1, is implemented on a general-purpose processor. A block diagram of the GACT array is shown in Figure 5a. It consists of a systolic array of processing elements (inspired by [44]), several SRAM banks that store the trace-back pointers, and trace-back logic.

**Figure 5:**
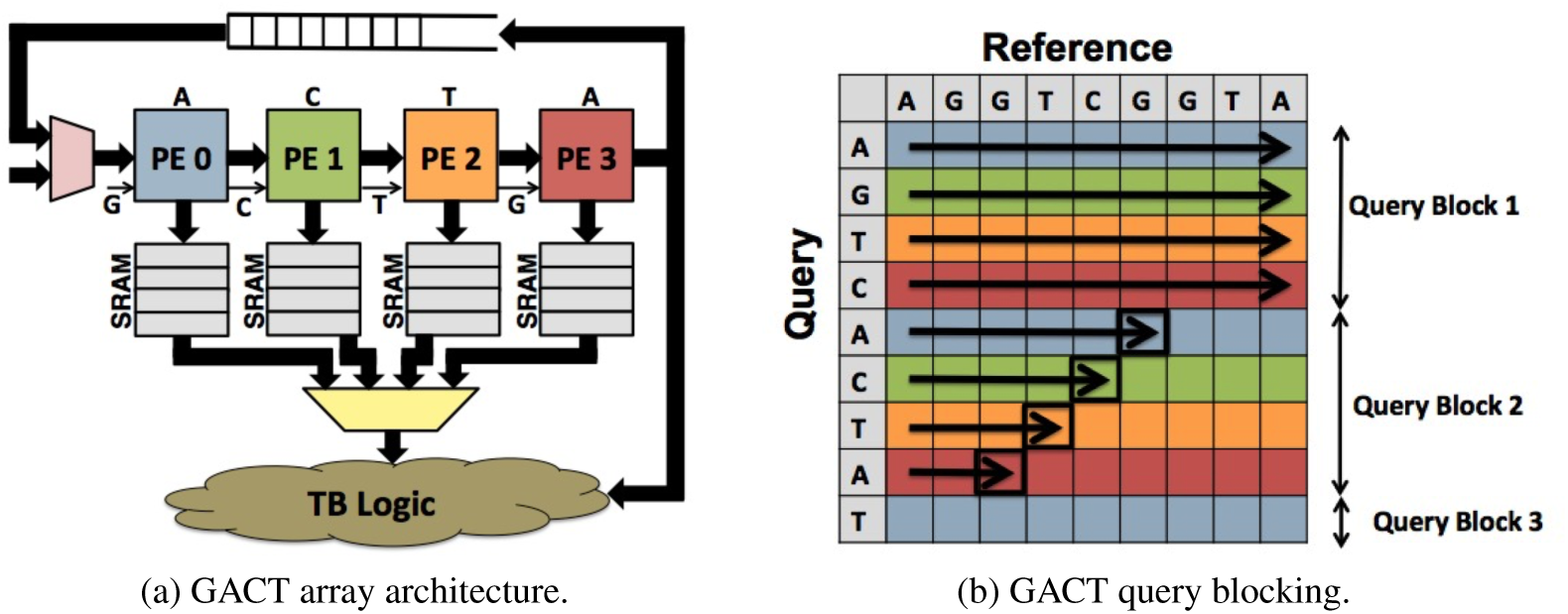
GACT hardware design. (a) Systolic architecture of GACT array with *N*_*pe*_=4. (b) Query blocking for *T*=9.

### Input/Output interface

The host processor provides to the GACT array: (i) a set of scoring parameters for the alignment, (ii) a tile size *T*, (iii) a reference (*R*^*tile*^) and a query (*Q*^*tile*^) sequence corresponding to the tile being processed, and (iv) a single bit *t* to indicate whether the trace-back should start from the highest scoring cell in the tile, or from the bottom-right cell of the tile. We have implemented GACT for the Smith-Waterman algorithm with affine gap penalties [45], and therefore, there are 18 scoring parameters - 16 parameters for the substitution matrix *W*, one parameter for gap open penalty *o* and one parameter for gap extend penalty *e*. The 18 parameters are stored using 10-bit signed integers in a register file and can be reused across several alignments. The maximum tile size is *T*_*max*_ = 512, because of the amount of trace-back memory provisioned in hardware (128KB). Our GACT implementation accepts ASCII inputs for *R*^*tile*^ and *Q*^*tile*^, but internally stores the sequences using 3-bits for an extended DNA alphabet Σ_*ext*_ = {*A,C, G, T, N*}, where *N* represents an unknown nucleotide and does not contribute to the alignment score.

Depending on whether *t* is set (unset), the GACT array outputs (i) score of the highest-scoring (bottom-right) cell of the tile, (ii) the coordinate (*i*^*∗*^, *j*^*∗*^) of the highest-scoring (bottom-right) cell of the tile, and (iii) a series of trace-back pointers for the optimal alignment in the tile from the highest-scoring (bottom-right) cell, where each pointer requires 2-bit to indicate whether the next operation in trace-back is an insert, a delete or a match.

### Processing Element (PE)

An array of *N*_*pe*_ PEs exploits *wavefront parallelism* to compute the *N*_*pe*_ cells of the DP-matrix each cycle. Each clock cycle, each PE computes three scores and one trace-back pointer for one cell of the DP-matrix. The three scores are:

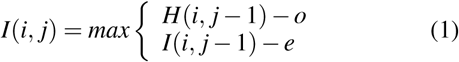

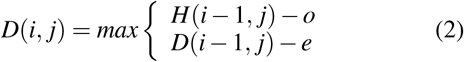

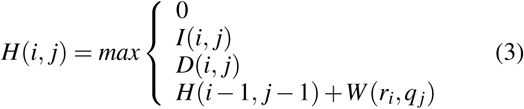

A four-bit trace-back pointer is also computed: 1 bit each for equations 1 and 2 to indicate whether the insertion and deletion gap to the cell resulted from opening or extending another gap, and 2 bits for equation 3 pointing to whether the cell’s best score results from a null (terminating), horizontal, vertical or diagonal cell. When *t* is set, each PE maintains the maximum score and corresponding position for the cells it has computed. On completion, the global maximum score and position is computed in a systolic fashion.

Figure 5a shows an example GACT array with *N*_*pe*_ = 4 PEs computing the DP-matrix of Figure 5b, that corresponds to a single GACT tile with *T* = 9. For *T* > *N*_*pe*_, the rows of the DP-matrix are divided into a number of query blocks, where a block consists of up to *N*_*pe*_ rows, one row for each PE. During the processing of one query block, each PE is initialized with the query base-pair corresponding to its row in the block. The reference sequence is then streamed through the array, shifting one base-pair every cycle. The systolic array architecture helps exploit the wave-front parallelism available in the computation of the query block. A FIFO is used to store the *H* and *D* scores of equations 3 and 2 of the last PE of the array, which gets consumed by the first PE during the computation of the next query block. The depth of this FIFO is *T*_*max*_, corresponding to the maximum number of columns in the DP-matrix.

### Trace-back Memory

A total memory of size 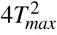 bits is required to store the trace-back pointers corresponding to the tile of size *T*_*max*_. In GACT array, this memory is divided into *N*_*pe*_ independent single-port SRAM banks, one back for every PE as shown in Figure 5a. Each cycle, each PE writes 4-bits for the trace-back pointer corresponding to the cell it computed, starting from address 0 and incrementing the address each cycle. As a result, pointers corresponding to the rows of consecutive query blocks get stored sequentially.

### Trace-back Logic

The trace-back (TB) logic is a finite-state machine (FSM) that traces back the optimal alignment in the tile starting from the cell provided to it as input. This FSM requires three cycles to produce 2-bits, which indicates whether the next direction in the trace-back is a match, insert, delete or terminate. in the first cycle, TB logic computes the address and the PE id corresponding to the next cell in the trace-back path depending on the direction pointer of the current cell. The address and PE id of the starting cell of the trace-back is provided by the last PE of the GACT array, which, depending on *t*, corresponds to either the highest-scoring or the bottom-right cell. This address is broadcast to all the banks of trace-back memory. The second cycle is consumed for the data to be available at the output of each SRAM bank. The third cycle is used by a multiplexer, which uses the incoming data from all SRAM banks and selects the one corresponding to the PE id provided by the TB logic. The output of the multiplexer is also used by the TB logic to generate the next 2-bit output direction in the same cycle. The trace-back terminates when either a terminating pointer is encountered or an edge of the DP-matrix (first row or column) is reached. The TB logic also keeps track of the number of base-pairs of *R*^*tile*^ and *Q*^*tile*^ in the alignment, which are used as *i*_*off*_ and *j*_*off*_ in Algorithm 1.

### Tile processing time

Using the proposed architecture, the approximate number of cycles *C*_*T*_ to process one tile of size *T* is given by:

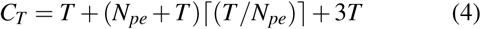

The first term of the equation 4 corresponds to the total cycles consumed in initializing the PEs, one cycle for each base-pair in the *Q*^*tile*^. The second term corresponds to the number of cycles required to process one query block (*N*_*pe*_ + *T*), times the number of query blocks, 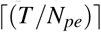. Finally, the last term corresponds to the number of cycles required by TB logic to compute trace-back path, assuming *T* directions in the alignment path. Equation 4 has been verified to be approximately correct using RTL simulation with genomic sequences.

In Darwin, we have used *N*_*pe*_ = 64, and therefore, for *T* = 300, roughly 3K cycles are required to processing the tile. Darwin uses *T*_*max*_ = 512, which requires 128KB SRAM for trace-back memory, with each of the 64 PEs connected to one 2KB SRAM bank.

## 7. EXPERIMENTAL METHODOLOGY

### Reference genome and read data

We used the latest human genome assembly GRCh38 as the reference genome for reference-guided assembly and our experiments involving GACT evaluation. We used only the haplotype chromosomes (chromosome 1-22, X, Y) and removed the mitochondrial genome, unmapped and unlocalized contigs from the assembly. The resulting assembly size is 3.08Gbp. The long reads for different technologies were simulated using PBSIM [46]. With simulated reads, ground truth is known, which allows for accurate measurements of sensitivity and specificity. We generate three sets of reads with length of 10Kbp each using a 30× coverage of the GRCh38 assembly: to match the error profiles of PacBio, high-quality Oxford Nanopore 2D reads and low quality Oxford Nanopore 1D reads, respectively. Default settings of continuous long read (CLR) model were used in PBSIM for PacBio reads at an error rate of 15.0%, while the PBSIM settings for ONT reads were taken from [29]. Table 1 shows the error profile for the three sets of reads. These parameters correspond roughly to the error profiles observed using P6-C4 chemistry for PacBio reads, and R7.3 chemistry for Oxford Nanopore reads. For *de novo* assembly, we used PBSIM to generate PacBio reads for 30× coverage of WS220 assembly of the *C. elegans* genome, resulting in 3.0Gbp of raw reads.

**Table 1:** Error profile of the reads sets. A 2.0% standard deviation was used for each read set.

| Read type | Substitution | Insertion | Deletion | Total |
| --- | --- | --- | --- | --- |
| PacBio | 1.50% | 9.02% | 4.49% | 15.01% |
| ONT 2D | 16.50% | 5.10% | 8.40% | 30.0% |
| ONT 1D | 20.39% | 4.39% | 15.20% | 39.98% |

### Hardware setup

Darwin, and the comparison baselines were run on an Intel Xeon E5-26200 CPU operating at 2.0GHz with 64GB DDR-3 memory. Hardware-acceleration of GACT was implemented on a PicoComputing platform consisting of six M505 modules on an EX-500 backplane. Each module contains one Xilinx Kintex 7 FGPA [47]. The FPGA is clocked at 250MHz and the communication with host CPU uses × 16 PCIe 2.0. For fair comparison, we used a single software thread for baseline techniques (discussed below) and a single software thread with only one GACT array for Darwin. The software thread was kept idle for Darwin while the FPGA was computing, and vice versa.

### Comparison baseline

For reference-guided assembly of PacBio reads, Darwin was compared to BWA-MEM [1] (version 0.7.14) using its -`pacbio` flag. For Oxford Nanopore reads, we used GraphMap [29], a recently developed and highly-sensitive mapper optimized for ONT reads. We note that while BWA-MEM also has a mode optimized for 2D reads of Oxford Nanopore, its sensitivity on our ONT_2D read dataset was only 89.1%, much lower than 98.1% that we observed using GraphMap. We used DALIGNER [2] as a baseline technique for *de novo* assembly of long reads. We tuned D-SOFT’s parameters to match or exceed the sensitivity of the baseline technique. We define sensitivity and specificity as:

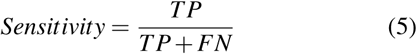

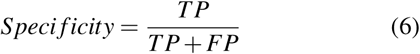

where TP, FP and FN are the number of true positives, false positives and false negatives of the algorithm, respectively. For reference-guided assembly, a true positive is when the read gets aligned to the reference within 50bp of the region of the ground-truth alignment reported by PBSIM. For *de novo* assembly, an overlap between two reads is considered a true positive if the two reads overlap by at least 1Kbp according to PBSIM, and if at least 80% of that overlap is detected by the algorithm under consideration. To evaluate the filtration ability of D-SOFT, we have also defined false positive rate (FPR) as the average number of false positives for every true positive resulting after D-SOFT filtration, without using GACT.

We also compared performance of GACT to other heuristics, namely X-drop [36] and Banded Smith-Waterman algorithm [35]. Both heuristics were implemented using SeqAn [48], a highly-optimized software library for sequence alignment. The alignments were carried out on the simulated reads and the reference genome and the performance is measured in terms of the ratio of alignment score to that of optimal Smith-Waterman algorithm, also implemented using SeqAn.

### ASIC Synthesis, Layout and Performance

To estimate the performance achievable with an application-specific integrated circuit implementation of GACT (rather than an FPGA implementation), we synthesized one processing element (PE), and one GACT array (including trace-back logic) with *N*_*pe*_ = 64 using Synopsys Design Compiler (DC) in a TSMC 40nm CMOS process. We used the tcbn40lpbwp standard cell library with worst case PVT corner. Placement and routing were carried out using Synopsys IC compiler (ICC) with 7 metal layers. We used Cacti [49] to estimate the area and power of the trace-back SRAM memory. We used the synthesis power estimation provided by Synopsys Design Compiler to estimate logic power. We compare the ASIC power to software power measured using Intel’s PCM power utility [50]. Since the utility only provides power at package level, and not core level, we estimate the application’s CPU power to be the difference of package power when application is running and when it is not.

To compute ASIC performance, we first computed the runtime component of GACT using Darwin’s FPGA prototype. This was done by executing every application twice on the FPGA, once with, and once without GACT. The difference in the two runtimes was estimated to be the runtime component of D-SOFT. We also created a log of the number of bytes exchanged between the host processor and the FPGA over PCIe, and estimated the communication overhead by measuring runtime of a dummy application exchanging same number of bytes between host and FPGA. The ASIC runtime was estimated to be the sum of D-SOFT runtime and the GACT runtime on FPGA, scaled to ASIC frequency. We assume that the ASIC implementation would have a fast communication interface with host processor such that GACT would be compute-bound. To better understand Darwin’s software bottlenecks, we used Gprof [51].

## 8. RESULTS AND DISCUSSION

### GACT

Figure 6 shows worst-case GACT score loss (in %) compared to optimal alignment using Smith-Waterman with a simple scoring scheme (match = +1, mismatch = -1, gap = -1) for different tile sizes and similarity of the sequences being aligned. The plot has been generated for 1,000 randomly chosen 10Kbp reads generated using PBSIM for sequence similarities of 70%, 80%, and 90%. For a given tile size, score loss for GACT decreases as the sequence similarity increases. For 100% similarity (not shown), GACT performs as well as optimal - perfectly tracing back along the main diagonal. The worst-case loss for GACT is lower than 0.95% for tile sizes above 300, and there is diminishing return for higher tile size, so we set *T*=300 for both, reference-guided and *de novo* assembly.

**Figure 6:**
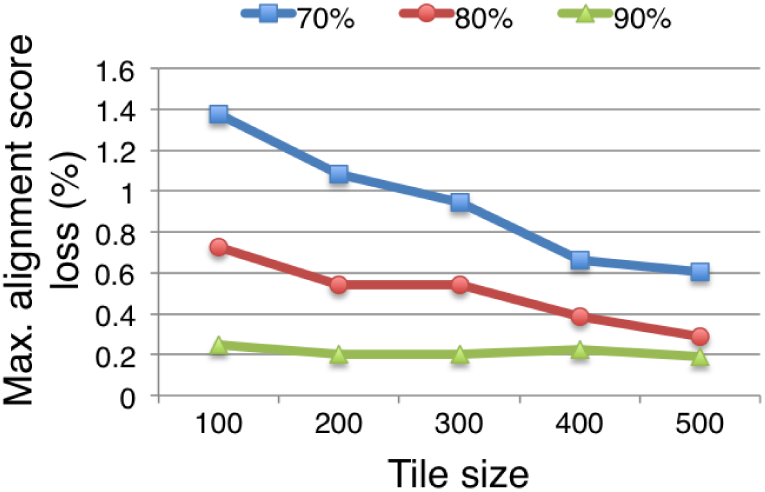
Maximum alignment score loss (%) using GACT for different sequence similarity and tile sizes.

Table 2 compares GACT to other alignment heuristics, namely Banded Smith-Waterman and X-drop for different trace-back memory requirements. We use a random sample of 10,000 10 Kbp sequences of PacBio read set in Table 1 and use simple scoring (match: +1, mismatch: -1, gap: -1) for the alignment. Since 4 bits are needed to store each trace-back pointer, we set the band *B* and threshold *X* in Banded Smith-Waterman and X-drop heuristic to *M/*(0.5*L*), where *M* is the memory size in bytes and *L* is the read length (10Kbp). Tile size *T* of GACT for *M*B of trace-back memory is 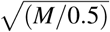, and is independent of the read length *L*. Banded Smith-Waterman, which has been implemented in numerous hardware-accelerators [21, 52, 22], does not work for long read alignment because the insertion rate is 4.53% higher than the deletion rate in PacBio reads (Table 1). This causes the optimal alignment to shift an average of 453bp from the main diagonal over a 10Kbp read. To capture the optimal alignment would require a band width of at least 453. Since X-drop adapts the band while computing the trace-back path,it has a far better score loss at 100KB memory and above. With only 50KB of memory, however, GACT has only 0.27% worst-case score loss, compared to 99.9% in X-drop. GACT is better suited for implementation on a memory-limited ASIC than current heuristics for long read alignments. Moreover, GACT performance is independent of the length of reads aligned, as opposed to the heuristics, whose performance would drop with a constant memory as read lengths increase.

**Table 2:** Average and max score loss of GACT compared to Banded Smith-Waterman and X-Drop heuristics on PacBio reads for different trace-back memory.

| Trace-back<br>memory (KB) | Mean score loss |  |  | Max score loss |  |  |
| --- | --- | --- | --- | --- | --- | --- |
|  | Banded | X-Drop | GACT | Banded | X-Drop | GACT |
| 20 | 98.7% | 99.51 | 0.14% | 99.8% | 99.9% | 0.31% |
| 50 | 95.2% | 6.8% | 0.10% | 98.9% | 99.9% | 0.27% |
| 100 | 89.2% | 0.13% | 0.03% | 94.35% | 0.19% | 0.19% |
| 1000 | 57.5% | 0.0% | 0.0023% | 67.2% | 0.0% | 0.06% |

Figure 7 shows why Banded Smith-Waterman performs much worse than GACT on long read alignments. The figure shows the DP-matrix for the two heuristics aligning a 1Kbp region from a PacBio read to the reference genome (GRCh38). The gray regions denotes the cells of the DP-matrix computed by the two heuristics, corresponding to *B*=25bp in Banded Smith-Waterman and *T*=220bp in GACT. The optimal trace-back path is shown in red. *B* and *T* were chosen such that the two heuristics had the same memory requirement for this alignment. PacBio reads have slightly higher insertion rate than deletion rate (Table 1), and we can see that the optimal trace-path path drifts from the main diagonal by 41bp. As a result, a band *B*=25bp can only capture a small portion of the optimal alignment that stays within the band shown in gray, as its alignment score is only 47% of the optimal. On the other hand, successive tiling in GACT in Figure 7 has been able to follow the optimal trace-back path perfectly, and in this case, GACT also resulted in the optimal alignment. Also note that the overlap between successive tiles is small, and so the number of tiles in an alignment of length *L* can be approximated as *L/T*.

**Figure 7:**
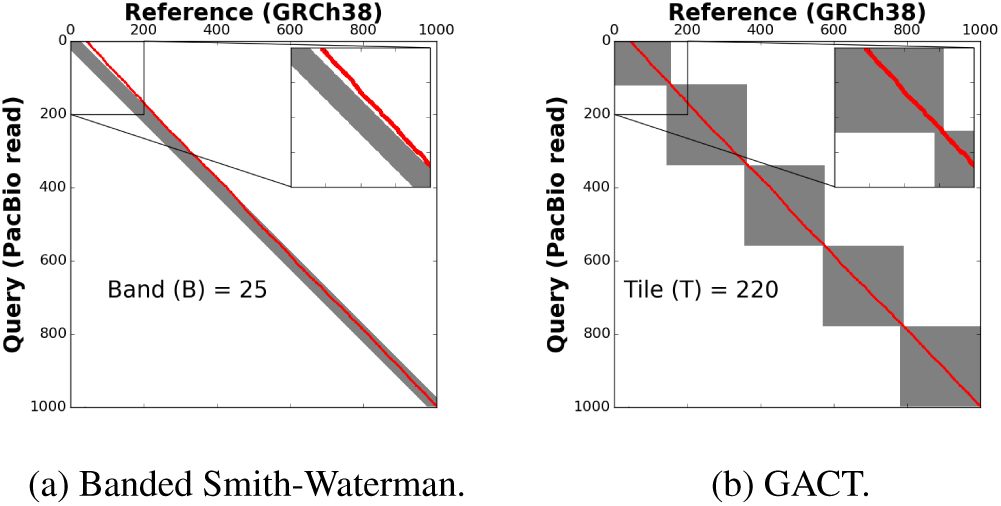
Comparison of DP-matrix of Banded Smith-Waterman and GACT for aligning 1Kbp PacBio read to reference (GRCh38). Gray cells of the matrix represent the cells computed by each algorithm. Red path represents optimal trace-back.

Figure 8 shows the layout of a single PE of the GACT array in a TSMC 40nm CMOS process. The PE fits in an area 55*um* on each side, 0.003*mm*^2^. A 64PE GACT array (excluding TB memory) requires 0.27*mm*^2^, the additional area is for control, trace-back logic, and the FIFO used for storing scores between query blocks. The critical path has a delay of 1.18ns, enabling operation at 847MHz (3.4× the FPGA frequency). The array consumes 17mW of power. A 128KB TB memory, for *T*_*max*_ = 512, requires another 1.10*mm*^2^ and consumes 120mW of additional power. Therefore, one GACT array has a total area of 1.37*mm*^2^ and power of 137mW.

**Figure 8:**
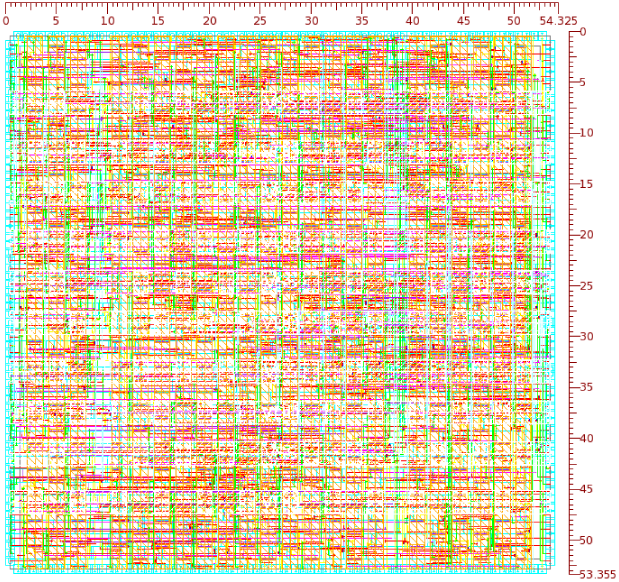
Layout of one GACT Processing Element (PE) in a TSMC 40nm CMOS process. Distances are in *µ*m.

Figure 9 shows the throughput of the hardware-accelerated GACT algorithm using *T*=300 for pairwise alignment of sequences with different lengths. The plot shows that GACT can perform 23,562 pairwise alignments of sequences of 1Kbp length every second. This throughput is roughly 35% of the expected throughput from equation 4 using 847MHz clock frequency and 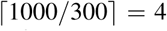 tiles, since 65% of the time is spent in software to extract successive tiles after each alignment and prepending/appending trace-back pointers (Algorithm 1). Even at this throughput, GACT hardware is 762× faster than highly-optimized software implementation using SeqAn, which could perform only 31 pairwise alignments using Banded SmithWaterman with *B*=45. Moreover, SeqAn’s software consumed 7W of power, making it 39,000× less energy-efficient than the GACT accelerator. Figure 9 also illustrates that the GACT throughput indeed scales inversely to the length of the alignment — the throughput reduced by nearly 10×, from 23,562 to 2,350 alignments per second when the length of the alignment increased by a factor of 10, from 1Kbp to 10Kbp. In comparison, throughput of the optimal algorithm would scale inverse of the square of sequence length. With this throughput, an oracular filter, which knows the read to reference genome mapping beforehand, would complete the reference-guided assembly of 54× human genome in only 1.96 hours with one GACT array, and the performance scales linearly with the number of arrays.

**Figure 9:**
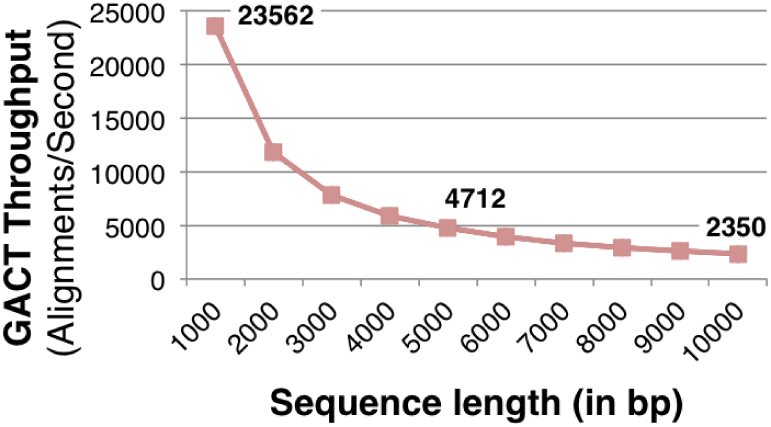
GACT throughput vs. sequence length (*T*=300). Plot generated using 10,000 sequences for every sequence length.

### D-SOFT

Figure 10 shows the sensitivity and false positive rate (FPR) of the D-SOFT filter (without dynamic programming filter) on PacBio reads. It highlights the one key advantage of the D-SOFT - it’s ability to filter out the false positives with negligible loss in sensitivity. This is achieved using the knob of the base-count threshold (*h*). In figure 10, with *h*=15, the filter can find a true positive in 99.87% cases, but it also results in 9079.2 false positives for every true positive. Increasing *h* to 24 decreases the sensitivity by only 0.16%, but decreases the FPR by 121×. Increasing *h* further to 30 decreases FPR marginally by another 1.86×, with additional 0.02% loss of sensitivity. The ability to be orders of magnitude more specific with little loss of sensitivity makes D-SOFT advantageous for Darwin, as it can now employ highly-sensitive, hardware-accelerated GACT for further filtering. Besides *h*, *k* and *N* can also be tuned in D-SOFT for different sensitivity and FPR trade-off. Higher FPR results in lower throughput, as false positives require further filtration using GACT.

**Figure 10:**
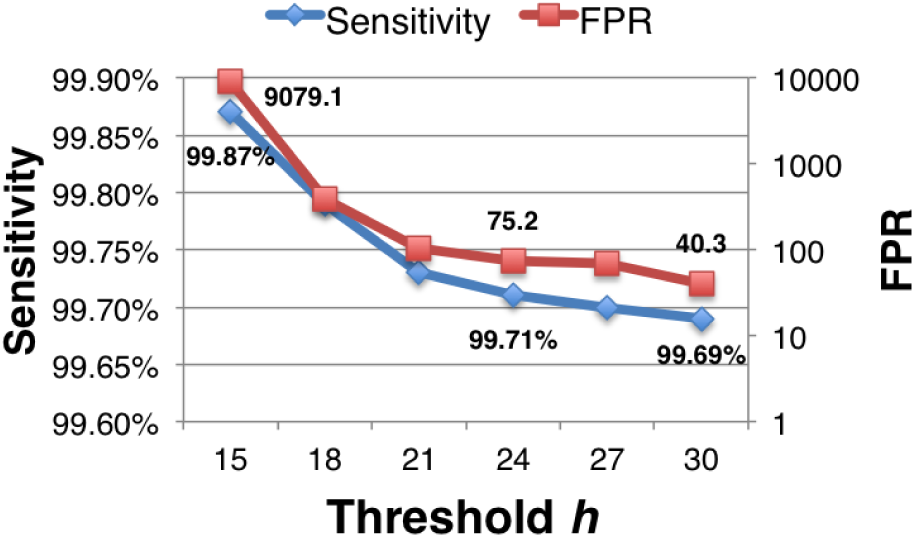
Sensitivity and false positive rate (FPR) for D-SOFT on PacBio reads for different choice of threshold *h*. Rest of D-SOFT settings: *k*=14, *N*=750.

### Darwin

Table 3 compares Darwin’s performance with baseline algorithms for reference-guided assembly on human genome using 50,000 randomly sampled reads for each of the three read sets of Table 1. We used *T*=300 and *h*_*tile*_=70 for GACT. We adjusted the D-SOFT parameters in Darwin, listed in Table 3, to match or exceed the sensitivity of the baseline technique. We note that for higher read error rate, smaller seed size (*k*) and larger number of seeds (*N*) are required to maintain high sensitivity. For PacBio reads, we observed a 125× speedup over the baseline technique, BWA-MEM, with 3.76% extra sensitivity. The extra sensitivity of Darwin can be explained, in part, by the choice of fixed length seeds of length 14 in Darwin, compared to supermaximal exact match (SMEM) seeding approach, as well as the additional chain filtering stage used in BWA-MEM, both of which discard some true positives. For ONT 2D and 1D reads, Darwin’s speedup is 107× and 12.47× that of GraphMap, respectively. Darwin uses very small seeds (*k*=11) for matching the high sensitivity of GraphMap in case of ONT 1D reads, and this resulted in D-SOFT slowdown, due to large number of seed hits, as well as GACT slowdown, due to alignments on too many false positives resulting from D-SOFT. GraphMap itself uses seeds with *k*=12, but it still maintains higher sensitivity because of two reasons: (i) it uses spaced seeds, which are known to provide higher sensitivity [40], and (ii) it uses multiple seeds with different shapes for every position in the query. We believe that incorporating the two aforementioned GraphMap strategies in D-SOFT could potentially speed up Darwin further, but we have left this evaluation for our future work. Darwin was also found to be more specific than baseline techniques, and this is because we discarded reads that resulted in less than 80% base-pairs aligning to the reference genome, and this resulted in fewer number of mapped reads.

**Table 3:** Comparison of Darwin and baseline techniques on reference-guided assembly.

| Read type | D-SOFT settings<br>($k, N, h$) | Mapped reads (%) | | Sensitivity | | Specificity | | Throughput (reads/sec) | |
| --- | --- | --- | --- | --- | --- | --- | --- | --- | --- |
|  |  | Baseline | Darwin | Baseline | Darwin | Baseline | Darwin | Baseline | Darwin |
| PacBio | (14, 750, 24) | 99.996% | 99.786% | 95.95% | 99.71% | 95.95% | 99.91% | 3.45 | 431.7 |
| ONT 2D | (12, 1000, 22) | 98.99% | 98.81% | 98.11% | 98.4% | 99.1% | 99.62% | 0.19 | 20.47 |
| ONT 1D | (11, 1300, 25) | 98.82% | 98.33% | 97.10% | 97.40% | 98.5% | 99.22% | 0.17 | 2.12 |

Figure 11 shows the histogram of scores observed for the first GACT tile for the true and the false positives resulting from D-SOFT for the three read sets. False positives from all three read sets are combined into a single histogram. The figure shows that PacBio reads, which have lower error rates, result in higher average GACT tile score. The figure also shows that it is easy to filter out a majority D-SOFT’s false positives using a simple threshold on first GACT tile score itself. With *h*_*tile*_=70 used in Table 3, 95.9% of D-SOFT’s false positives can be filtered out after the first GACT tile itself, with less than 0.05% additional loss in sensitivity. Because Darwin uses dynamic programming as a filter following the initial seeding using D-SOFT, it retains the sensitivity of D-SOFT almost exactly. In contrast, nearly all competing long read aligners, including BWA-MEM, BLASR, and GraphMap, use several filtration stages with each stage incorporating a heuristic having a non-trivial influence on sensitivity.

**Figure 11:**
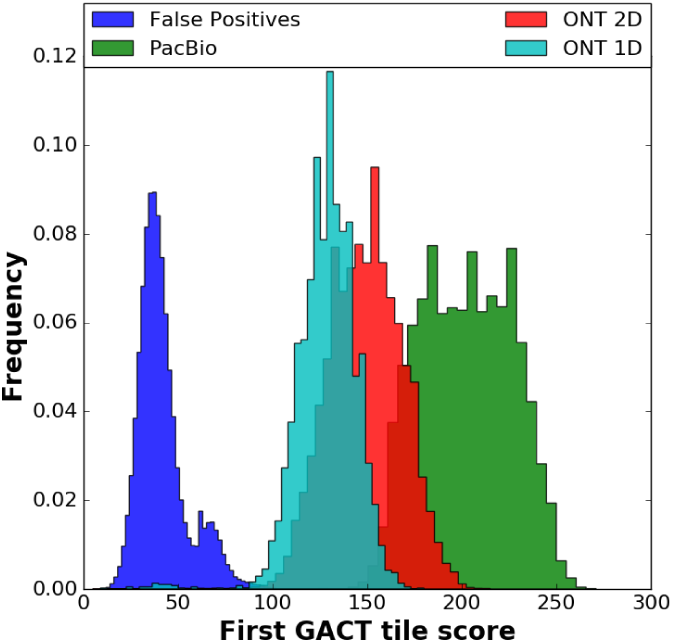
Histogram of first GACT tile score for D-SOFT false positives and true positives of different read sets.

Figure 12 shows the fraction of mismatching base-pairs between the consensus sequences generated using GACT and Smith-Waterman algorithm. The figure is generated by first mapping all reads from chromosome 1 (consisting of 247Mbp) in the PacBio read set of Table 1 using GACT and optimal Smith-Waterman algorithm for alignment, and then for each alignment algorithm, considering the consensus of first *C* reads covering a base-pair, to find the consensus of that base-pair at a coverage of *C*. We find that at a low coverage *C*=1, nearly 1bp in every 1100bp has a mismatch between GACT and optimal (Smith-Waterman) consensus, but this fraction drops exponentially, and at 13× or higher coverage, we observe a perfect match of consensus sequence using GACT and optimal alignment. We believe this result is because GACT introduces deviations from the optimal alignment at a small number of random locations, and the stochastic nature of the deviations implies that they could be corrected statistically, much like read errors can be corrected using consensus of several reads. However, read errors occur much more frequently, and hence need a much higher coverage to be corrected for. [17] observed error rate of 0.7% persists at even 15× coverage in PacBio reads. The result in Figure 12 indicates that with high coverage sequencing, Darwin can be used reliably for applications in precision medicine, such as *variant calling* in patients [53].

**Figure 12:**
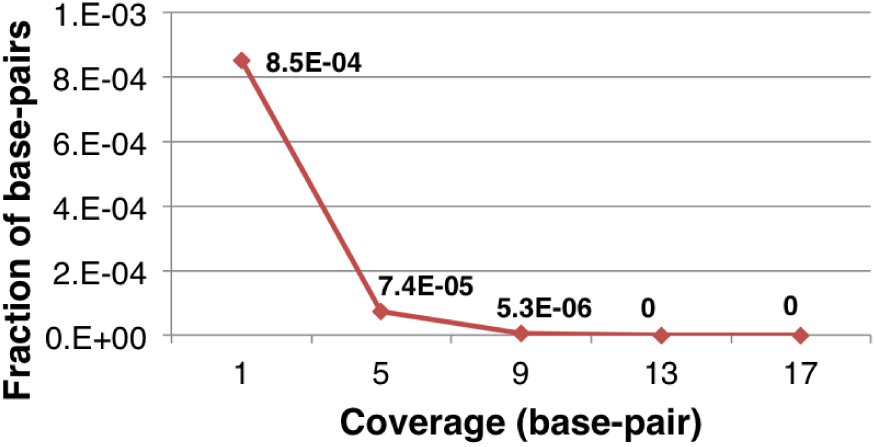
Fraction of mismatching base-pairs between the consensus sequences generated using GACT and Smith-Waterman algorithm.

Table 4 shows the comparison of Darwin and DALIGNER for finding pairwise overlaps between PacBio reads for *de novo* assembly of a 30× *C. elegans* genome. DALIGNER detected 99.8% overlaps in over 14 hours, and in comparison, Darwin required less than one hour to find 99.89% overlaps, a 15.6× speedup over DALIGNER. Of this, 99.1% overlaps were directly derived from D-SOFT, and remaining 0.79% overlaps were transitively determined. 14 min of Darwin’s 55 min were required to build the seed lookup table. Although DALIGNER has a more rigid filter requiring at least 35bp to be conserved in a diagonal band using seeds of size 14bp, compared to at least 24bp conservation in D-SOFT settings for the same seed size, DALIGNER was found to be more sensitive. This is because (i) DALIGNER uses all seeds of the reads, whereas Darwin uses only *N*=1100 seeds from the head of every read, (ii) Darwin discards high-frequency seeds while constructing the seed lookup table, but DALIGNER maintains all seeds.

**Table 4:** Comparison of DALIGNER and Darwin on overlap assembly of 30× *C. elegans* genome. D-SOFT settings are (*k*=14, *N*=1100, *h*=24).

|  | Sensitivity | Specificity | Runtime |
| --- | --- | --- | --- |
| DALIGNER | 99.80% | 88.3% | 14h21m |
| Darwin | 99.89% | 90.1% | 0h55m |

### Darwin’s bottleneck

Profiling of Darwin revealed that GACT contributes to nearly 20% of the overall runtime, including the time spent on filtration of false positives resulting from D-SOFT, while D-SOFT itself contributes to about 80%. This shows that 762× speedup obtained by ASIC was critical to Darwin’s overall speedup, but also that scaling up GACT arrays further would provide incremental benefits. D-SOFT is the current bottleneck in Darwin, and detailed profiling using Gprof revealed that around 70-90% of D-SOFT runtime is spent updating the two arrays, *last*_*hit*_*pos* and *bp*_*count* (line 10-11 in Algorithm 2), but *SeedLookup* is fast. This is because *SeedLookup* has majority sequential memory accesses, compared to completely random memory accesses in updating the two arrays. Speeding up of random memory accesses using specialized hardware is possible, as shown in recent work using e-DRAM for Graph Analytics workloads [54], and extending this approach to D-SOFT could help further scale Darwin’s performance. Only 120MB memory is required for the two arrays making it possible to place this entire memory on-chip.

## 9. RELATED WORK

### Alignment heuristics and hardware-acceleration

GACT algorithm falls under the rich body of literature involving heuristic approximation to the Smith-Waterman algorithm for pairwise alignment of sequences. Navarro et. al [55] have provided a detailed survey of algorithms used for approximate string matching, with applications ranging from genome sequence analysis to text retrieval. Besides X-drop and Banded Smith-Waterman, a technique proposed by Gene Myers [56] that aligns two strings using dynamic programming, while exploiting bit-vector parallelism available in hardware, has found immense traction in genomics applications. However, unlike GACT, X-drop and Banded Smith-Waterman, [56] restricts the number of mismatches between the two sequences to be at most *k*, where *k* is a parameter to the algorithm. Moreover, unlike GACT, [56] has a memory requirement that grows linearly with the length of sequences being aligned. In fact, to the best of our knowledge, GACT is the first approximation algorithm to Smith-Waterman that takes constant trace-back memory requirement into consideration, which makes it suitable for hardware acceleration, especially for aligning long sequences.

Acceleration of sequence alignment has also been well explored in hardware architecture and compiler communities. Lipton et. al [44] were the first to propose a systolic array exploiting wavefront parallelism to perform Smith-Waterman alignment of arbitrarily long sequences. This architecture, however did not handle trace-back. A number of papers [57, 58, 23, 24] have implemented variants of systolic arrays in hardware, particularly on FPGAs. While they have all achieved high speedup against software, they only handle the matrix fill step of Smith-Waterman, leaving the trace-back step to software, which would require the software to recompute the score matrix around high-scoring cells. This undermines the benefits of hardware acceleration, particularly in the context of read assembly. Some prior work [59, 60] has proposed hardware-acceleration of read mapping for second generation technologies, without trace-back in hardware. Compiler assisted high level synthesis of systolic arrays has been studied in [61, 62]. This work is orthogonal to GACT, but GACT hardware synthesis could potentially improve from the proposed techniques.

Some more recent work [22, 21, 63, 64] has also implemented the trace-back step in hardware. Chen et. al [21] accelerated long read mapping using Banded Smith-Waterman with a maximum read length of 4Kbp. Reads are now longer in length and as our results indicate, Banded Smith-Waterman does not work well with different insertion and deletion rates observed in current sequencing technologies. Nawaz et. al [22] also accelerated banded Smith-Waterman in hardware, for sequences up to 10Kbp in length, but their architecture required 2.5MB of trace-back memory. In comparison, the proposed GACT architecture requires only 128KB, and can align arbitrarily long sequences, even with different insertion and deletion rates. We consider this to be a significant contribution for hardware-acceleration of sequence alignment, since all prior work on hardware acceleration has found limited scope of application due to the restriction of length of sequences that can be aligned. To our knowledge, GACT is the first hardware-acceleration technique to eliminate this restriction.

### Filtration heuristics

D-SOFT falls under the broad category of filtration heuristics that reduce the search space for dynamic programming alignment. In particular, D-SOFT was designed to be more specific than the filtration techniques that are based on counting the number of seed hits conserved in a band of diagonals. The first filtration stage of BLAST [19], two-hit BLAST [38], GraphMap [29], and BLASR [39] are all based on seed hit count. DALIGNER [2] was the first to our knowledge to directly count the base-pairs instead of seeds in a diagonal band, which allows it to be highly specific even at high sensitivity. This algorithm inspired D-SOFT. However, in implementation, D-SOFT and DALIGNER differ in significant ways. First, for the same memory constraint, DALIGNER creates more blocks because it stores the read number, the seed, and the offset of the seed for every seed in the block. D-SOFT, which uses a seed position table for the concatenated reads, stores a single offset. The seed, which translates to an address, is not stored. More read blocks requires comparing more block pairs. Second, DALIGNER uses a sort and merge operation for every block pair, while D-SOFT queries seeds only from the head of every read to detect overlaps with a read block. This results in high speedup, since fewer seeds are used when accessing the seed position table and this access is highly sequential. Finally, D-SOFT can be applied to reference-guided as well as overlap-based in *de novo* assembly. On the other hand, DALIGNER is specialized for the overlap step, and its extension to reference-guided assembly is non-trivial (reference genomes are several orders longer than individual reads).

There are filtration techniques, such as BWA-MEM [1], that are neither based on counting seeds in a band of diagonals nor the number of bases. Instead, BWA-MEM uses super maximal seeds and its performance has been evaluated in this paper. Other examples include Canu [65], M-HAP [66] and LSH-ALL-PAIRS [67], which are based on probabilistic, locality-sensitive hashing of seeds. While D-SOFT cannot generalize to these techniques, as our results on other filtration techniques indicate, it is still possible to tune D-SOFT parameters to match or exceed the sensitivity of these heuristics.

Finally, there has been work on improving the sensitivity of seed-based filtration and reducing the storage requirement of seeds. These are orthogonal to D-SOFT. Work on spaced seeds [40], transition-constrained seeds [68] and vector seeds [69], help improve seeding sensitivity. Minimizers [70] help reduce the storage requirement of seeds.

### Sequence alignment frameworks

Like Darwin, some prior work has focused on implementation of a complete sequence alignment framework, implementing filtration as well as sequence alignment with relatively flexible parameters, and aided by hardware acceleration. Examples include TimeLogic [71], which has implemented FPGA-based framework for BLAST and HMMER, and Edico Genome [64], which provides FPGA-based framework for acceleration of BWA-GATK pipeline for reference-guided assembly on second generation sequencing. Darwin leverages the tunable sensitivity of D-SOFT, and can accelerate alignment of arbitrarily long sequences using GACT, overcoming two major limitations of prior frameworks. In this paper, we have shown that Darwin handles and provides high speedup versus hand-optimized software for two distinct applications: reference-guided and *de novo* assembly of reads, and can work with reads with very different error rates. To our knowledge, Darwin is the first hardware-accelerated framework to demonstrate speedup in more than one class of applications and in future, we plan to extend Darwin to alignment applications even beyond read assembly.

## 10. CONCLUSION AND FUTURE WORK

*Darwin* is a hardware-accelerated framework for genomic sequence alignment that is orders of magnitude faster than software counterparts used in long read assembly. Darwin uses (i) D-SOFT, a novel filtration heuristic, that has tunable sensitivity and is highly specific even at high sensitivity for assembling noisy long reads, and (ii) GACT, a novel alignment algorithm based on dynamic programming that can be used for aligning arbitrarily long sequences and accelerated by orders of magnitude using constant memory hardware. Darwin has been shown to give large speedup for reference-guided as well as *de novo* assembly of long reads, with different error profiles.

We plan to extend Darwin by building an accelerator for D-SOFT, the current bottleneck in Darwin. This, combined with scaling up number of GACT arrays, is likely to add another order of magnitude speedup. We also plan to evaluate spaced seeds and seeding with multiple shapes to further improve D-SOFT sensitivity. Several extant species will be sequenced over the next decade, and we plan to extend Darwin to applications beyond read assembly, such as whole genome alignments and remote homology search, both of which will contribute profoundly to genomic science as more genome sequences become available.

## 11. ACKNOWLEDGMENTS

We sincerely thank Professor Serafim Batzoglou for his valuable insights into long read sequencing. We thank Albert Ng, who generously shared the RTL code of his Smith-Waterman accelerator. We thank Professor Boris Murmann who provided the access to ASIC CAD tools. We also thank NVIDIA, Xilinx and the Platform Lab (Stanford University) for supporting this research, through funding and equipment donation. We thank all members of CVA Lab and Bejerano Lab for providing valuable feedback.

## REFERENCES

[1] H. Li, “Aligning sequence reads, clone sequences and assembly contigs with bwa-mem,” arXiv preprint arXiv:1303.3997, 2013.

[2] G. Myers, “Efficient local alignment discovery amongst noisy long reads,” in International Workshop on Algorithms in Bioinformatics, pp. 52–67, Springer, 2014.

[3] E. S. Lander, L. M. Linton, B. Birren, C. Nusbaum, M. C. Zody, J. Baldwin, K. Devon, K. Dewar, M. Doyle, W. FitzHugh, et al., “Initial sequencing and analysis of the human genome,” Nature, 2001.

[4] J. C. Venter, M. D. Adams, E. W. Myers, P. W. Li, R. J. Mural, G. G. Sutton, H. O. Smith, M. Yandell, C. A. Evans, R. A. Holt, et al., “The sequence of the human genome,” Science, 2001.

[5] Z. D. Stephens, S. Y. Lee, F. Faghri, R. H. Campbell, C. Zhai, M. J. Efron, R. Iyer, M. C. Schatz, S. Sinha, and G. E. Robinson, “Big data: astronomical or genomical?,” PLoS Biol, 2015.

[6] “Illumina HiSeq X Series of Sequencing Systems: Specification Sheet., http://www.illumina.com/documents/products/datasheets/datasheet-hiseq-x-ten.pdf, Accessed: 2016-11-04.”

[7] M. Eisenstein, “Oxford nanopore announcement sets sequencing sector abuzz,” Nature biotechnology, vol. 30, no. 4, pp. 295–296, 2012.

[8] M. A. Hamburg and F. S. Collins, “The path to personalized medicine,” New England Journal of Medicine, vol. 363, no. 4, pp. 301–304, 2010.

[9] E. D. Pleasance, R. K. Cheetham, P. J. Stephens, D. J. McBride, S. J. Humphray, C. D. Greenman, I. Varela, M.-L. Lin, G. R. Ordóñez, G. R. Bignell, et al., “A comprehensive catalogue of somatic mutations from a human cancer genome,” Nature, vol. 463, no. 7278, pp. 191–196, 2010.

[10] N. Krumm, T. N. Turner, C. Baker, L. Vives, K. Mohajeri, K. Witherspoon, A. Raja, B. P. Coe, H. A. Stessman, Z.-X. He, et al., “Excess of rare, inherited truncating mutations in autism,” Nature genetics, vol. 47, no. 6, pp. 582–588, 2015.

[11] C. López-Otín, M. A. Blasco, L. Partridge, M. Serrano, and G. Kroemer, “The hallmarks of aging,” Cell, vol. 153, no. 6, pp. 1194–1217, 2013.

[12] O. Spichenok, Z. M. Budimlija, A. A. Mitchell, A. Jenny, L. Kovacevic, D. Marjanovic, T. Caragine, M. Prinz, and E. Wurmbach, “Prediction of eye and skin color in diverse populations using seven SNPs,” Forensic Science International: Genetics, vol. 5, no. 5, pp. 472–478, 2011.

[13] J. Z. Li, D. M. Absher, H. Tang, A. M. Southwick, A. M. Casto, S. Ramachandran, H. M. Cann, G. S. Barsh, M. Feldman, L. L. Cavalli-Sforza, et al., “Worldwide human relationships inferred from genome-wide patterns of variation,” science, vol. 319, no. 5866, pp. 1100–1104, 2008.

[14] T. C. Sequencing, A. Consortium, et al., “Initial sequence of the chimpanzee genome and comparison with the human genome,” Nature, vol. 437, no. 7055, pp. 69–87, 2005.

[15] E. E. Schadt, S. Turner, and A. Kasarskis, “A window into third-generation sequencing,” Human molecular genetics, vol. 19, no. R2, pp. R227–R240, 2010.

[16] D. Gordon, J. Huddleston, M. J. Chaisson, C. M. Hill, Z. N. Kronenberg, K. M. Munson, M. Malig, A. Raja, I. Fiddes, L. W. Hillier, et al., “Long-read sequence assembly of the gorilla genome,” Science, vol. 352, no. 6281, p. aae0344, 2016.

[17] J. Eid, A. Fehr, J. Gray, K. Luong, J. Lyle, G. Otto, P. Peluso, D. Rank, P. Baybayan, B. Bettman, et al., “Real-time dna sequencing from single polymerase molecules,” Science, vol. 323, no. 5910, pp. 133–138, 2009.

[18] S. Goodwin, J. Gurtowski, S. Ethe-Sayers, P. Deshpande, M. C. Schatz, and W. R. McCombie, “Oxford nanopore sequencing, hybrid error correction, and de novo assembly of a eukaryotic genome,” Genome research, vol. 25, no. 11, pp. 1750–1756, 2015.

[19] S. F. Altschul, W. Gish, W. Miller, E. W. Myers, and D. J. Lipman, “Basic local alignment search tool,” Journal of molecular biology, 1990.

[20] R. S. Harris, Improved pairwise alignment of genomic DNA. ProQuest, 2007.

[21] P. Chen, C. Wang, X. Li, and X. Zhou, “Accelerating the next generation long read mapping with the fpga-based system,” IEEE/ACM Transactions on Computational Biology and Bioinformatics (TCBB), vol. 11, no. 5, pp. 840–852, 2014.

[22] Z. Nawaz, M. Nadeem, H. van Someren, and K. Bertels, “A parallel fpga design of the smith-waterman traceback,” in Field-Programmable Technology (FPT), 2010 International Conference on, pp. 454–459, IEEE, 2010.

[23] P. Faes, B. Minnaert, M. Christiaens, E. Bonnet, Y. Saeys, D. Stroobandt, and Y. Van de Peer, “Scalable hardware accelerator for comparing dna and protein sequences,” in Proceedings of the 1st international conference on Scalable information systems, p. 33, ACM, 2006.

[24] X. Jiang, X. Liu, L. Xu, P. Zhang, and N. Sun, “A reconfigurable accelerator for smith–waterman algorithm,” IEEE Transactions on Circuits and Systems II: Express Briefs, vol. 54, no. 12, pp. 1077–1081, 2007.

[25] T. F. Smith and M. S. Waterman, “Identification of common molecular subsequences,” Journal of molecular biology, vol. 147, no. 1, pp. 195–197, 1981.

[26] W. J. Kent, “BLAT - the BLAST-like alignment tool,” Genome research, 2002.

[27] H. Li and R. Durbin, “Fast and accurate long-read alignment with burrows–wheeler transform,” Bioinformatics, vol. 26, no. 5, pp. 589–595, 2010.

[28] R. Li, C. Yu, Y. Li, T.-W. Lam, S.-M. Yiu, K. Kristiansen, and J. Wang, “Soap2: an improved ultrafast tool for short read alignment,” Bioinformatics, vol. 25, no. 15, pp. 1966–1967, 2009.

[29] I. Sović, M. Šikić, A. Wilm, S. N. Fenlon, S. Chen, and N. Nagarajan, “Fast and sensitive mapping of nanopore sequencing reads with graphmap,” Nature communications, vol. 7, 2016.

[30] S. Koren and A. M. Phillippy, “One chromosome, one contig: complete microbial genomes from long-read sequencing and assembly,” Current opinion in microbiology, vol. 23, pp. 110–120, 2015.

[31] Z. Li, Y. Chen, D. Mu, J. Yuan, Y. Shi, H. Zhang, J. Gan, N. Li, X. Hu, B. Liu, et al., “Comparison of the two major classes of assembly algorithms: overlap–layout–consensus and de-bruijn-graph,” Briefings in functional genomics, vol. 11, no. 1, pp. 25–37, 2012.

[32] G. A. Auwera, M. O. Carneiro, C. Hartl, R. Poplin, G. del Angel, A. Levy-Moonshine, T. Jordan, K. Shakir, D. Roazen, J. Thibault, et al., “From fastq data to high-confidence variant calls: the genome analysis toolkit best practices pipeline,” Current protocols in bioinformatics, pp. 11–10, 2013.

[33] X. Chen, O. Schulz-Trieglaff, R. Shaw, B. Barnes, F. Schlesinger, M. Källberg, A. J. Cox, S. Kruglyak, and C. T. Saunders, “Manta: rapid detection of structural variants and indels for germline and cancer sequencing applications,” Bioinformatics, vol. 32, no. 8, pp. 1220–1222, 2016.

[34] D. S. Hirschberg, “A linear space algorithm for computing maximal common subsequences,” Communications of the ACM, vol. 18, no. 6, pp. 341–343, 1975.

[35] K.-M. Chao, W. R. Pearson, and W. Miller, “Aligning two sequences within a specified diagonal band,” Computer applications in the biosciences: CABIOS, vol. 8, no. 5, pp. 481–487, 1992.

[36] Z. Zhang, S. Schwartz, L. Wagner, and W. Miller, “A greedy algorithm for aligning dna sequences,” Journal of Computational biology, vol. 7, no. 1-2, pp. 203–214, 2000.

[37] C. L. Ip, M. Loose, J. R. Tyson, M. de Cesare, B. L. Brown, M. Jain, R. M. Leggett, D. A. Eccles, V. Zalunin, J. M. Urban, et al., “Minion analysis and reference consortium: Phase 1 data release and analysis,” F1000Research, vol. 4, 2015.

[38] S. F. Altschul, T. L. Madden, A. A. Schäffer, J. Zhang, Z. Zhang, W. Miller, and D. J. Lipman, “Gapped blast and psi-blast: a new generation of protein database search programs,” Nucleic acids research, vol. 25, no. 17, pp. 3389–3402, 1997.

[39] M. J. Chaisson and G. Tesler, “Mapping single molecule sequencing reads using basic local alignment with successive refinement (blasr): application and theory,” BMC bioinformatics, vol. 13, no. 1, p. 238, 2012.

[40] U. Keich, M. Li, B. Ma, and J. Tromp, “On spaced seeds for similarity search,” Discrete Applied Mathematics, vol. 138, no. 3, pp. 253–263, 2004.

[41] H. Xin, S. Nahar, R. Zhu, J. Emmons, G. Pekhimenko, C. Kingsford, C. Alkan, and O. Mutlu, “Optimal seed solver: optimizing seed selection in read mapping,” Bioinformatics, vol. 32, no. 11, pp. 1632–1642, 2016.

[42] D. Gussfield, “Algorithms on strings, trees, and sequences,” Computer Science and Computional Biology (Cambrigde, 1999), 1997.

[43] T. W. Lam, W.-K. Sung, S.-L. Tam, C.-K. Wong, and S.-M. Yiu, “Compressed indexing and local alignment of dna,” Bioinformatics, vol. 24, no. 6, pp. 791–797, 2008.

[44] R. J. Lipton and D. P. Lopresti, Comparing long strings on a short systolic array. Princeton University, Department of Computer Science, 1986.

[45] O. Gotoh, “An improved algorithm for matching biological sequences,” Journal of molecular biology, vol. 162, no. 3, pp. 705–708, 1982.

[46] Y. Ono, K. Asai, and M. Hamada, “Pbsim: Pacbio reads simulatorâĂŤ toward accurate genome assembly,” Bioinformatics, vol. 29, no. 1, pp. 119–121, 2013.

[47] “Pico computing product brief: M-505-K325T, http://picocomputing.com/wp-content/uploads/2013/11/product-brief-m-505-k325t.pdf, Accessed: 2016-11-04.”

[48] A. Döring, D. Weese, T. Rausch, and K. Reinert, “Seqan an efficient, generic c++ library for sequence analysis,” BMC bioinformatics, vol. 9, no. 1, p. 11, 2008.

[49] P. Shivakumar and N. P. Jouppi, “Cacti 3.0: An integrated cache timing, power, and area model,” tech. rep., Technical Report 2001/2, Compaq Computer Corporation, 2001.

[50] “Intel PCM power utility., https://software.intel.com/en-us/articles/intel-performance-counter-monitor#pcm_power, Accessed: 2016-11-04.”

[51] S. L. Graham, P. B. Kessler, and M. K. Mckusick, “Gprof: A call graph execution profiler,” in ACM Sigplan Notices, vol. 17, pp. 120–126, ACM, 1982.

[52] “A banded Smith-Waterman FPGA accelerator for Mercury BLASTP, author=Harris, Brandon and Jacob, Arpith C and Lancaster, Joseph M and Buhler, Jeremy and Chamberlain, Roger D, booktitle=2007 International Conference on Field Programmable Logic and Applications, pages=765–769, year=2007, organization=IEEE,”

[53] H. Li, J. Ruan, and R. Durbin, “Mapping short dna sequencing reads and calling variants using mapping quality scores,” Genome research, vol. 18, no. 11, pp. 1851–1858, 2008.

[54] T. J. Ham, L. Wu, N. Sundaram, N. Satish, and M. Martonosi, “Graphicionado: A high-performance and energy-efficient accelerator for graph analytics,” in Microarchitecture (MICRO), 2016 49th Annual IEEE/ACM International Symposium on, pp. 1–13, IEEE, 2016.

[55] G. Navarro, “A guided tour to approximate string matching,” ACM computing surveys (CSUR), vol. 33, no. 1, pp. 31–88, 2001.

[56] G. Myers, “A fast bit-vector algorithm for approximate string matching based on dynamic programming,” Journal of the ACM (JACM), vol. 46, no. 3, pp. 395–415, 1999.

[57] Y. Yamaguchi, T. Maruyama, and A. Konagaya, “High speed homology search with fpgas,” in Proceedings of the 7th Pacific Symposium on Biocomputing (PSB’02), pp. 271–282, 2001.

[58] C. W. Yu, K. Kwong, K.-H. Lee, and P. H. W. Leong, “A smith-waterman systolic cell,” in New Algorithms, Architectures and Applications for Reconfigurable Computing, pp. 291–300, Springer, 2005.

[59] W. Tang, W. Wang, B. Duan, C. Zhang, G. Tan, P. Zhang, and N. Sun, “Accelerating millions of short reads mapping on a heterogeneous architecture with fpga accelerator,” in Field-Programmable Custom Computing Machines (FCCM), 2012 IEEE 20th Annual International Symposium on, pp. 184–187, IEEE, 2012.

[60] Y.-T. Chen, J. Cong, J. Lei, and P. Wei, “A novel high-throughput acceleration engine for read alignment,” in Field-Programmable Custom Computing Machines (FCCM), 2015 IEEE 23rd Annual International Symposium on, pp. 199–202, IEEE, 2015.

[61] D. Greaves, S. Sanyal, and S. Singh, “Synthesis of a parallel smith-waterman sequence alignment kernel into fpga hardware,”

[62] B. Buyukkurt and W. A. Najj, “Compiler generated systolic arrays for wavefront algorithm acceleration on fpgas,” in 2008 International Conference on Field Programmable Logic and Applications, pp. 655–658, IEEE, 2008.

[63] C. B. Olson, M. Kim, C. Clauson, B. Kogon, C. Ebeling, S. Hauck, and W. L. Ruzzo, “Hardware acceleration of short read mapping,” in Field-Programmable Custom Computing Machines (FCCM), 2012 IEEE 20th Annual International Symposium on, pp. 161–168, IEEE, 2012.

[64] P. Van Rooyen, R. J. McMillen, and M. Ruehle, “Bioinformatics systems, apparatuses, and methods executed on an integrated circuit processing platform,” Jan. 5 2016. US Patent App. 14/988,666.

[65] S. Koren, B. P. Walenz, K. Berlin, J. R. Miller, and A. M. Phillippy, “Canu: scalable and accurate long-read assembly via adaptive k-mer weighting and repeat separation,” bioRxiv, p. 071282, 2016.

[66] K. Berlin, S. Koren, C.-S. Chin, J. P. Drake, J. M. Landolin, and A. M. Phillippy, “Assembling large genomes with single-molecule sequencing and locality-sensitive hashing,” Nature biotechnology, vol. 33, no. 6, pp. 623–630, 2015.

[67] J. Buhler, “Efficient large-scale sequence comparison by locality-sensitive hashing,” Bioinformatics, vol. 17, no. 5, pp. 419–428, 2001.

[68] L. Noé and G. Kucherov, “Yass: enhancing the sensitivity of dna similarity search,” Nucleic acids research, vol. 33, no. suppl 2, pp. W540–W543, 2005.

[69] B. Brejová, D. G. Brown, and T. Vinař, “Vector seeds: An extension to spaced seeds,” Journal of Computer and System Sciences, vol. 70, no. 3, pp. 364–380, 2005.

[70] M. Roberts, W. Hayes, B. R. Hunt, S. M. Mount, and J. A. Yorke, “Reducing storage requirements for biological sequence comparison,” Bioinformatics, vol. 20, no. 18, pp. 3363–3369, 2004.

[71] “TimeLogic Corporation., http://www.timelogic.com, Accessed: 2016-11-04.”

